# Meta-analyses of the NEPTUNE dataset related to high and low risk FSGS and cellular APOL1 models identifies Calcium signalling, mTOR signaling and inflammation-associated pathways as operative in APOL1-mediated kidney disease

**DOI:** 10.64898/2026.07.30.741742

**Authors:** Wasco Wruck, Chantelle Thimm, Tama Amin, Maria-Darline Somoano Sanchez, Sejla Pulo, James Adjaye

## Abstract

**Background:** The variants G1 and G2 within the APOL1 gene confer a higher risk of APOL1-mediated kidney disease (AMKD) whilst associated with an evolutionary advantage against trypanosome-mediated sleeping sickness.

**Methods:** In this study, we analysed transcriptome data of kidney biopsies from FSGS patients with the APOL1 high-risk (HR) and low-risk (LR) variants and compared it to cellular models based on patient-specific podocytes, HEK cells with engineered over-expressing HR variants and HR variant single-cell-RNA-seq data from kidney organoids.

**Results:** We identified a signature of up- and down-regulated genes between biopsies from FSGS patients with APOL1 HR and LR variants. The up-regulated genes are functionally annotated to be associated with Calcium and mTOR signaling, whilst the down-regulated genes with inflammatory and immune response pathways. These pathways were confirmed by comparing with cellular models. Analysis of small molecules reverting the IFN-γ stimulated gene expression to the non-stimulated gene expression in genome-edited APOL-G1 kidney organoids revealed several putative candidates such as the mTOR inhibitor AZD-2014.

**Conclusion:** We have unveiled a signature of up- and down-regulated genes between APOL1 HR and LR kidney biopsies which could be assigned as associated with Calcium and mTOR signaling and down-regulated immune response.

## Introduction

The discovery of coding variants in apolipoprotein L1 (APOL1) has transformed our understanding of genetically mediated kidney disease. The APOL1 G1 (S342G and I384M) and G2 (N388/Y389) variants represent the strongest common genetic risk factors for non-diabetic CKD in individuals of recent African ancestry. Inheriting two APOL1 HR alleles markedly increases the risk of developing APOL1-mediated kidney disease (AMKD), including focal segmental glomerulosclerosis, HIV-associated nephropathy, hypertension-associated CKD, and accelerated progression to kidney failure [1]. Remarkably, these variants attained high population frequencies through positive evolutionary selection because they protect against African trypanosome infections. Whereas the ancestral APOL1 G0 protein lyses *trypanosoma brucei,* the G1 and G2 variants evade parasite-mediated neutralization, thereby conferring resistance to African sleeping sickness at the expense of increased susceptibility to kidney disease [2].

Although the genetic association between APOL1 and kidney disease is unequivocal, the mechanisms by which APOL1 HR variants cause renal injury remain incompletely resolved. Disease manifestation typically requires both inheritance of two APOL1 risk alleles and a pro-inflammatory “second hit”, such as viral infections (e.g., COVID-19 [3], HIV [4], which activate interferon signaling and induce APOL1 expression via the JAK/STAT signalling pathway [5], [6]. Once induced, APOL1 HR variants disrupt a broad spectrum of cellular processes that collectively contribute to kidney injury. A central pathogenic mechanism involves the formation of cation-permeable pores at the plasma membrane and intracellular organelles [7], [2] resulting in dysregulated potassium, sodium, and calcium fluxes that impair ion homeostasis and membrane integrity [2], [8]. These disturbances are closely linked to mitochondrial dysfunction [9], characterized by impaired oxidative phosphorylation, ATP depletion, excessive reactive oxygen species production, and activation of intrinsic apoptotic pathways [9]; [10]; [11]. In parallel, APOL1 risk variants induce endoplasmic reticulum stress [12], impair lysosomal function and autophagic flux [12], disrupt intracellular calcium homeostasis, inhibit protein translation, and activate multiple stress-responsive signaling cascades, including p38, JNK, MAPK8 and ERK [13]; [14]. These alterations promote inflammasome activation, inflammatory cytokine production, and several forms of regulated cell death, including apoptosis, pyroptosis, necroptosis, and ferroptosis [12], [15], [16].

Furthermore, APOL1-mediated cytoskeletal remodeling compromises podocyte architecture and slit diaphragm integrity [17]. Although these pathogenic pathways are well documented experimentally, it remains unclear whether they represent independent mechanisms or converge on a common upstream process that initiates APOL1-mediated cellular injury. Increasing evidence suggests that disruption of the APOL1 interactome may represent one such upstream mechanism.

The locations of the mutations driving the G1 and G2 variants within the binding domain of APOL1 to the trypanosomal Serum-Resistance-Antigen (SRA)- [18] may inform speculation about a similar human protein binding to APOL1 and this way preventing opening of ion channels and subsequent kidney impairment. The G1 and G2 variants would then inhibit or reduce binding of this human protein to the APOL1 SRA binding domain. While the SNARE protein VAMP8 might be a candidate [19],[20] Friedman and Pollak suggest that the APOL1 binding domain may be autoinhibitory [18]

Progress towards resolving these questions has been hindered by the lack of experimental models that faithfully recapitulate APOL1 biology. To date, most mechanistic studies have used immortalized cell lines, particularly HEK-derived systems, in which APOL1 risk variants are overexpressed. Although these models have provided important insights into APOL1-mediated pathogenic mechanisms, they frequently employ supraphysiological protein expression levels and fail to capture the cellular complexity and physiological microenvironment of the human kidney [21], [22], [23], [24], [25], [26], [27], [28], [29].

Patient-derived kidney biopsy samples provide clinically relevant material but are limited in availability and generally represent established disease, restricting their utility for studying early pathogenic events [30]. More recently, primary human renal cells, induced pluripotent stem cell (iPSC)-derived podocytes and endothelial cells [31]; [32] and kidney organoids have emerged as promising alternatives that more closely resemble human kidney biology [15]; [32]. Nevertheless, these systems remain technically challenging and are characterised by significant fluctuations and complexity. An overview of representative *in vitro* models in APOL1 research is provided in Table S1.

In this study, we first searched for differentially regulated genes and pathways between HR and LR in kidney biopsy-derived transcriptomes from NEPTUNE and compared them to cellular models based on human urine-derived podocyte-like epithelial cells (HUPEC) with the APOL1 G1/G2 variant, on HEK293 cells overexpressing the G1 and G2 variants and on kidney organoids with the G1 variant.

Employing over-representation analysis we identified functional annotations of the gene signature derived from the biopsy and its overlaps with the cellular models, thus condensing it to essential biological processes. These findings could help illuminate the molecular mechanisms underlying APOL1-mediated kidney injury.

## Methods

### Acquisition of kidney transcriptome data

Microarray datasets related to experiments conducted with kidney cells and podocytes were downloaded from the public repository at the NCBI GEO (National Center for Biotechnology Information, Gene Expression Omnibus) (Table 1). Figure S1 depicts a pipeline of the various analyses conducted on the datasets listed in Table 1.

**Table 1:** kidney transcriptome datasets from NCBI GEO used in this study.

| dataset | contents | kidney cell types | platform | PubMedId |
| --- | --- | --- | --- | --- |
| GSE135663 | G0/G1, IFN-γ | kidney organoids | Sc-RNAseq | 32656538 |
| GSE68125 | FSGS, MCD, MN, low / high risk<br>APOL1, IFN-γ | Glomerular biopsies | Affymetrix Human Gene 2.1 ST Array | 26150607 |
| GSE85918 | Tet-on G0, G1, G2 | HEK293 cells | Illumina HumanHT-12 V4.0 | 27821631 |
| GSE194330 | G0/G0,G1/G2, FSGS | Urine-derived podocyte epithelial cells (HUPECs) | RNAseq | 36644347 |

#### Data analysis

Transcriptome-based datasets pertaining to kidney biopsies (glomerular cells) and several in vitro kidney models derived from Affymetrix and Illumina microarray platforms and via RNAseq and single-cell-RNAseq were downloaded from NCBI GEO (Table 1). The glomerular biopsy dataset originated from the NEPTUNE study [30]. The in vitro model-based datasets were from Liu et al. [33], Ma et al. [9] and Yoshida et al. [34]. All datasets were fed into R/Bioconductor [35] for follow-up processing. Differentially expressed genes of the NEPTUNE glomerular dataset between the APOL1 HR and LR variants were determined by the moderated test statistics methods from the Bioconductor limma package [36] using a threshold of p < 0.05 and a log2-ratio > 0.415 for up-regulation and a log2-ratio < -0.415 for down-regulation. Subsequently, these up- and down-regulated gene signatures were employed to generate cluster analysis and gene expression heatmaps via the heatmap.2 function from the gplots package [37] in the NEPTUNE and the in vitro model datasets. For the cluster analysis complete linkage was used as an agglomeration method and Pearson correlation as similarity measure and colour bars indicating APOL1 variants were assigned. Venn diagrams of expressed genes were drawn via the R package VennDiagram [38].

#### Over-representation analysis of pathways and gene ontologies

Gene Ontologies (GOs) over-represented in the up- and down-regulated gene signatures associated with APOL1 HR were determined via the Bioconductor package GOstats [39]. Gene sets associated with KEGG and Reactome pathways downloaded from the KEGG database [40] in October 2025 and the Reactome database in May 2025 [41] were employed for over-representation analysis of the APOL1 HR up- and down-regulation gene-signatures via the R-built-in hypergeometric test. The results of KEGG, Reactome and GO analyses were visualized as dot plots showing p-values, ratios of significant genes and numbers of significant genes per pathway via the package ggplot2 [42]. The network of genes in enriched KEGG pathways was drawn via the Bioconductor packages enrichplot [43] and clusterprofiler [44] using the union of up-and down-regulation gene signatures.

#### Analysis of putative small molecules capable of reverting APOL1 HR-mediated gene expression

APOL1 G1 and G0 variant kidney organoids derived single-cell-RNAseq (sc-RNAseq) data were downloaded from the NCBI GEO accession GSE135663 and imported into the R environment via the R package Seurat [45]. sc-RNAseq data was filtered for cells with more than 200 features and mitochondrial gene percentage less than five percent and for genes within at least ten cells with at least three reads. The filtered data was processed via the standard Seurat pipeline and all cells were summarized for the G0 and G1 variants untreated or treated with IFN-γ. Up- and down-regulated genes between G1-IFN-γ and G0-IFN-γ were determined via the Seurat method FindMarkers using a p-value of 0.05 and a log2-fold-change of log2(1.5). The gene signatures of up- and down-regulated genes were uploaded to the web tool SigCom-LINCS [46] for analysis of small molecules (SMs). Via SigCom-LINCS the gene set is successively refined to the top regulated genes and to a cluster including the most consistently down-regulated genes.

## Results

### A signature of differentially expressed genes distinguishes between HR and LR APOL1 variants in glomerular cells derived from FSGS patient kidney biopsies

Figure 1 shows heatmaps of genes up- (Figure 1A) and down-regulated (Figure 1B) between HR and LR APOL1 variants in glomerular cells derived from patients with Focal Segmental Glomerusclerosis (FSGS, Suppl. Table S2). The signature of up-regulated genes as well as the signature of down-regulated genes (Suppl. Table S3) can segregate APOL1 HR and APOL1 LR transcriptomes.

**Figure 1:**
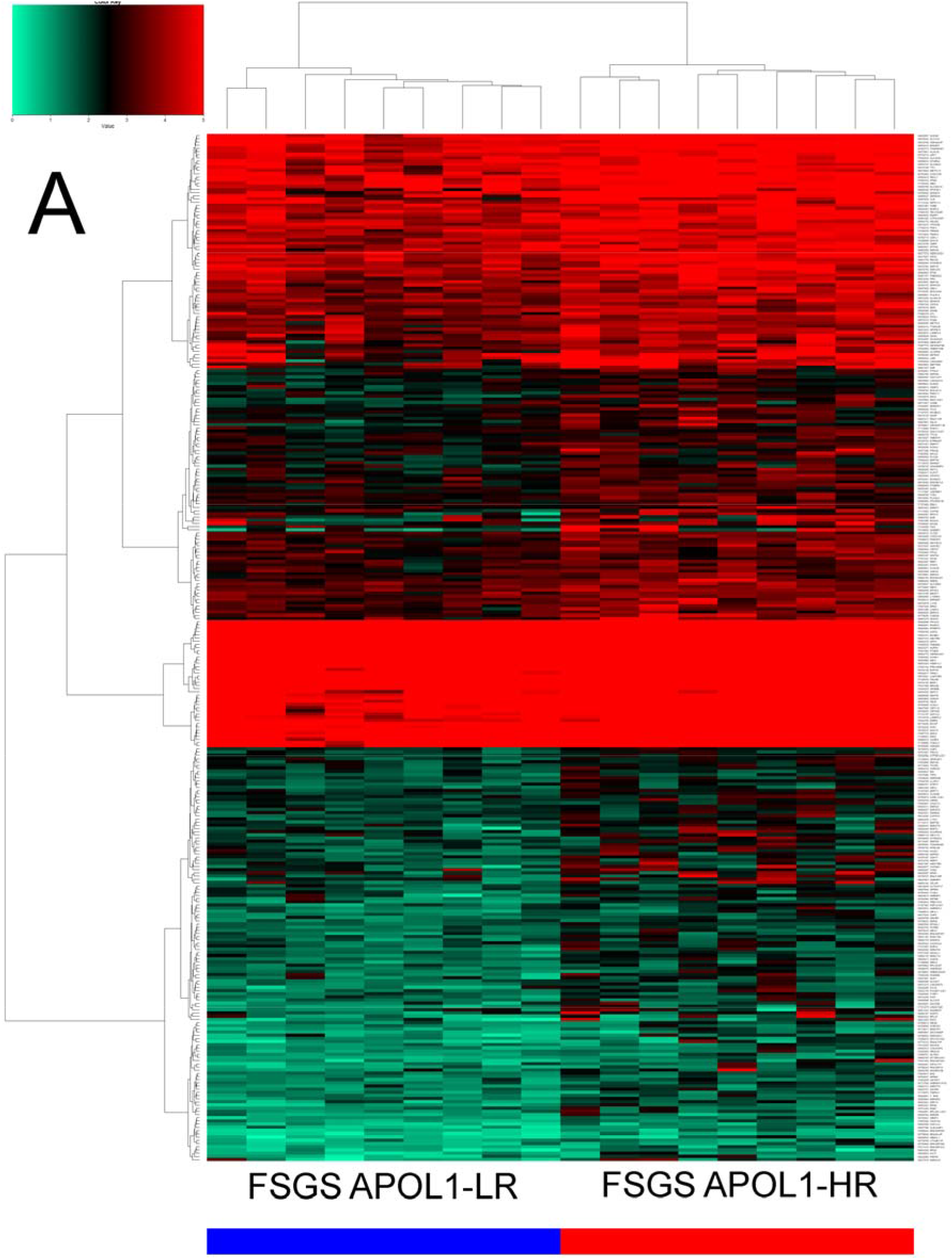
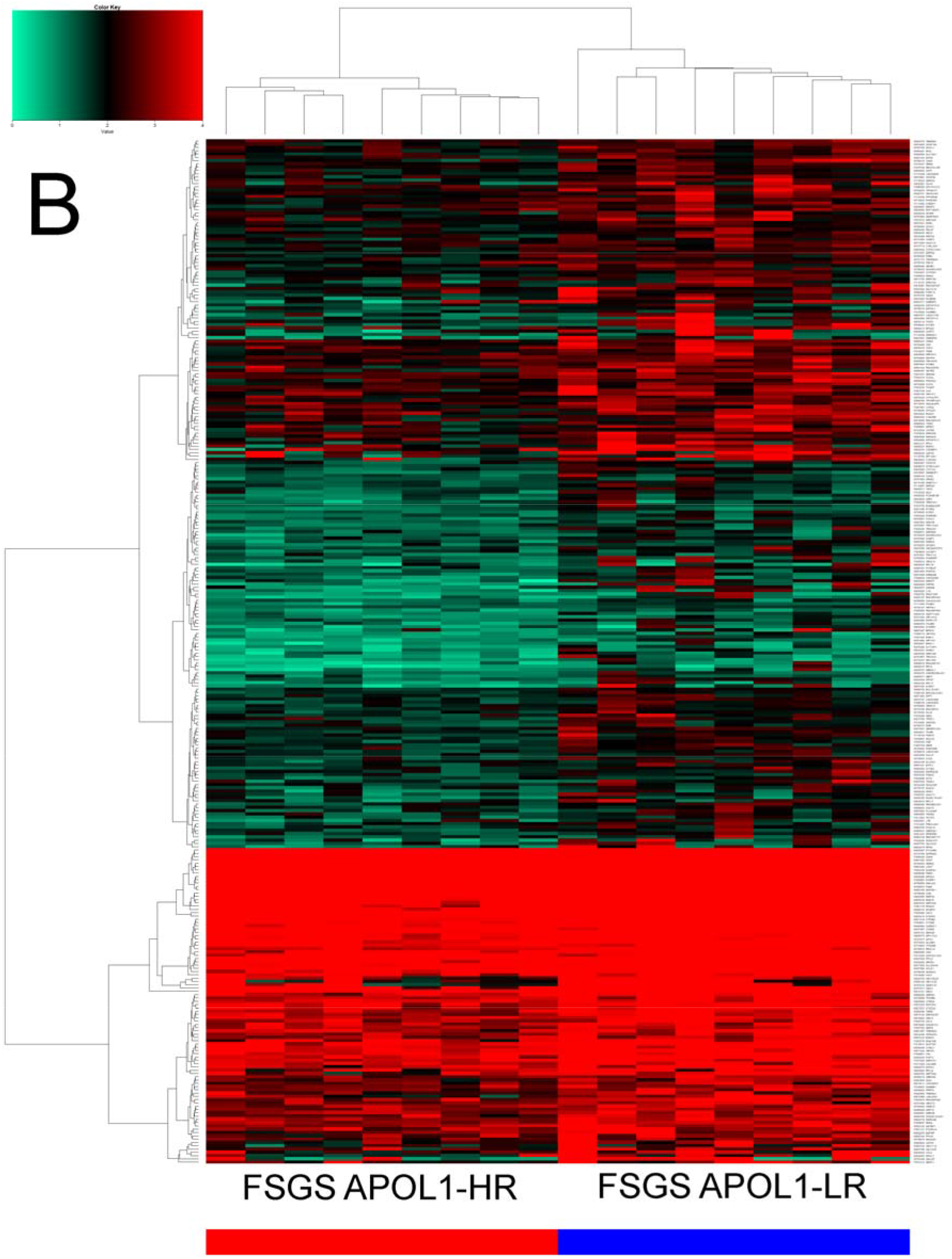
Gene signatures derived from glomerular cells can distinguish between FSGS patients with HR and LR APOL1 variants. (A) Genes up-regulated between APOL1 HR and LR, (B) genes down-regulated between APOL1 HR and APOL1 LR (color bar: red – APOL1 HR samples, blue – APOL1 LR samples).

### Differentially expressed genes between HR- and LR APOL1 variants in FSGS glomerular cells are associated with over-representation of Calcium and mTOR signaling

Figure 2 shows the results of a KEGG pathway analysis of genes up- and down-regulated between HR and LR APOL1 variants. The over-represented pathways in the up-regulated genes (Figure 2A, Table 2) include Calcium and mTOR signaling as well as neuron-associated pathways which we showed in an earlier publication to have considerable overlap with pathways active in podocytes [47]. The over-represented pathways in the down-regulated genes (Figure 2B) include inflammation-associated pathways such as Toll-like receptor and TNF signaling and also reactive oxygen species (ROS).

**Figure 2:**
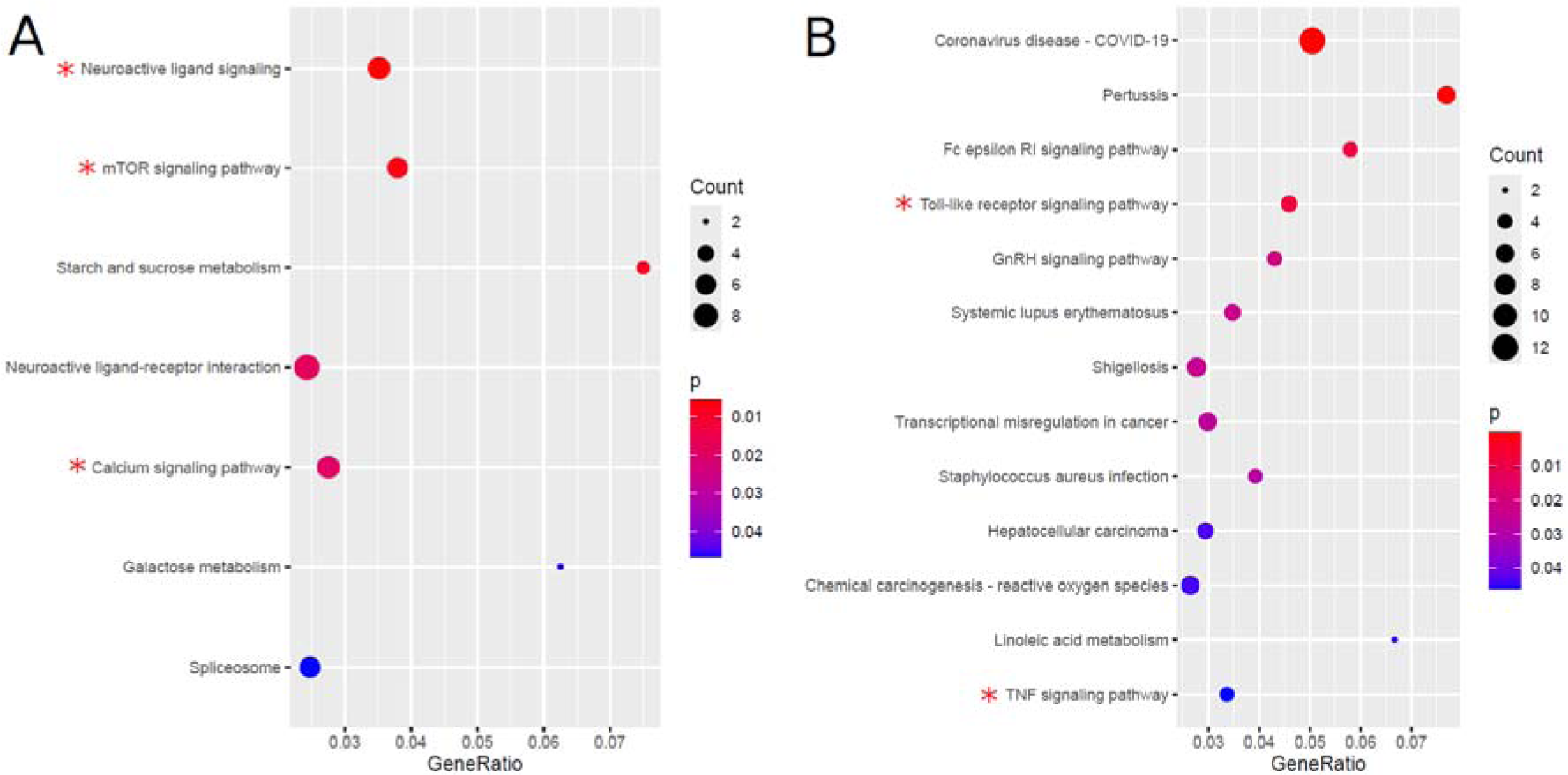
mTOR and Calcium signaling pathways are up-regulated between HR and LR APOL1 variants in patients with FSGS. (A) Over-represented pathways in genes up-regulated between HR and LR APOL1 variants include Neuroactive ligand signalling and ligand-receptor interactions, mTOR and Calcium signaling. (B) Over-represented pathways in genes down-regulated between HR and LR APOL1 variants include inflammation-related pathways such as Toll-like receptor and TNF signaling.

**Table 2:** KEGG pathway analysis results of genes up-regulated in APOL1-HR vs. LR.

| kegg_name | p | q | genes |
| --- | --- | --- | --- |
| Neuroactive ligand signaling | 0.0057 | 0.2561 | ADRA2C,GABBR1,GRIA3,OGFR,SLC5A7,TAC4,TACR2 |
| mTOR signaling pathway | 0.0072 | 0.2561 | CAB39L,LAMTOR3,RPS6,RRAGD,TTI1,WNT5A |
| Starch and sucrose metabolism | 0.0090 | 0.2561 | G6PC1,GAN C,GYG2 |
| Neuroactive ligand-receptor interaction | 0.0183 | 0.3373 | ADCYAP1,ADRA2C,GABBR1,GRIA3,OGFR,PTGER3,TAC4,TACR2,THRB |
| Calcium signaling pathway | 0.0198 | 0.3373 | EGF,NGF,PHKA1,PLCD4,PTGER3,TACR2,TNNC1 |
| Galactose metabolism | 0.0464 | 0.5267 | G6PC1,GAN C |
| Spliceosome | 0.0467 | 0.5267 | DDX39B,DHX16,HNRNPA1,HNRNPC,HNRNPK,PRPF8 |

**Table 3:** KEGG pathway analysis results of genes down-regulated in APOL1-HR vs. LR.

| kegg_name | p_hyper | q_hyper | genes |
| --- | --- | --- | --- |
| Coronavirus disease - COVID-19 | 0.00001 | 0.00210 | C4A,MAPK8,RPL11,RPL17,RPL19,RPL24,RPL4,RPL9,RPS10,RPS11,RPS24,RPS5 |
| Pertussis | 0.00024 | 0.01727 | C4A,CD14,CXCL5,IL23A,MAPK8,PYCARD |
| Fc epsilon RI signaling pathway | 0.00765 | 0.27738 | MAP2K1,MAP2K4,MAPK8,PLA2G4F |
| Toll-like receptor signaling pathway | 0.00781 | 0.27738 | CCL4,CD14,MAP2K1,MAP2K4,MAPK8 |
| GnRH signaling pathway | 0.02104 | 0.44767 | MAP2K1,MAP2K4,MAPK8,PLA2G4F |
| Systemic lupus erythematosus | 0.02363 | 0.44767 | C4A,H2AC11,H3-3A,H3C7,H4C3 |
| Shigellosis | 0.02459 | 0.44767 | CD14,H3-3A,H3C7,HCLS1,MAPK8,PYCARD,SEPTIN2 |
| Transcriptional misregulation in cancer | 0.02643 | 0.44767 | CD14,CDK9,DEFA1,H3-3A,H3C7,SUPT3H |
| Staphylococcus aureus infection | 0.02837 | 0.44767 | C4A,DEFA1,FCGR2B,KRT27 |
| Hepatocellular carcinoma | 0.04364 | 0.50154 | DPF1,FZD10,GSTM2,HGF,MAP2K1 |
| Chemical carcinogenesis - ROS | 0.04386 | 0.50154 | CYP1A2,GSTM2,HGF,MAP2K1,MAP2K4,MAPK8 |
| Linoleic acid metabolism | 0.04500 | 0.50154 | CYP1A2,PLA2G4F |
| TNF signaling pathway | 0.04592 | 0.50154 | CXCL5,MAP2K1,MAP2K4,MAPK8 |

### Crucial genes in the Calcium and mTOR pathway are up-regulated between the APOL1 HR and LR variants in FSGS patients

Figure 3 shows a pathway chart of Calcium signaling highlighting genes up-regulated in APOL1 HR variants in red and genes down-regulated in green. The pathway chart depicts that EGF, PTGER3, PLCD4, TNNC1 and PHKA1 are up-regulated in the HR variants resulting in contraction – potentially contraction of the glomerular tuft and vasoconstriction - while ORAI3 and TFEB are down-regulated.

**Figure 3:**
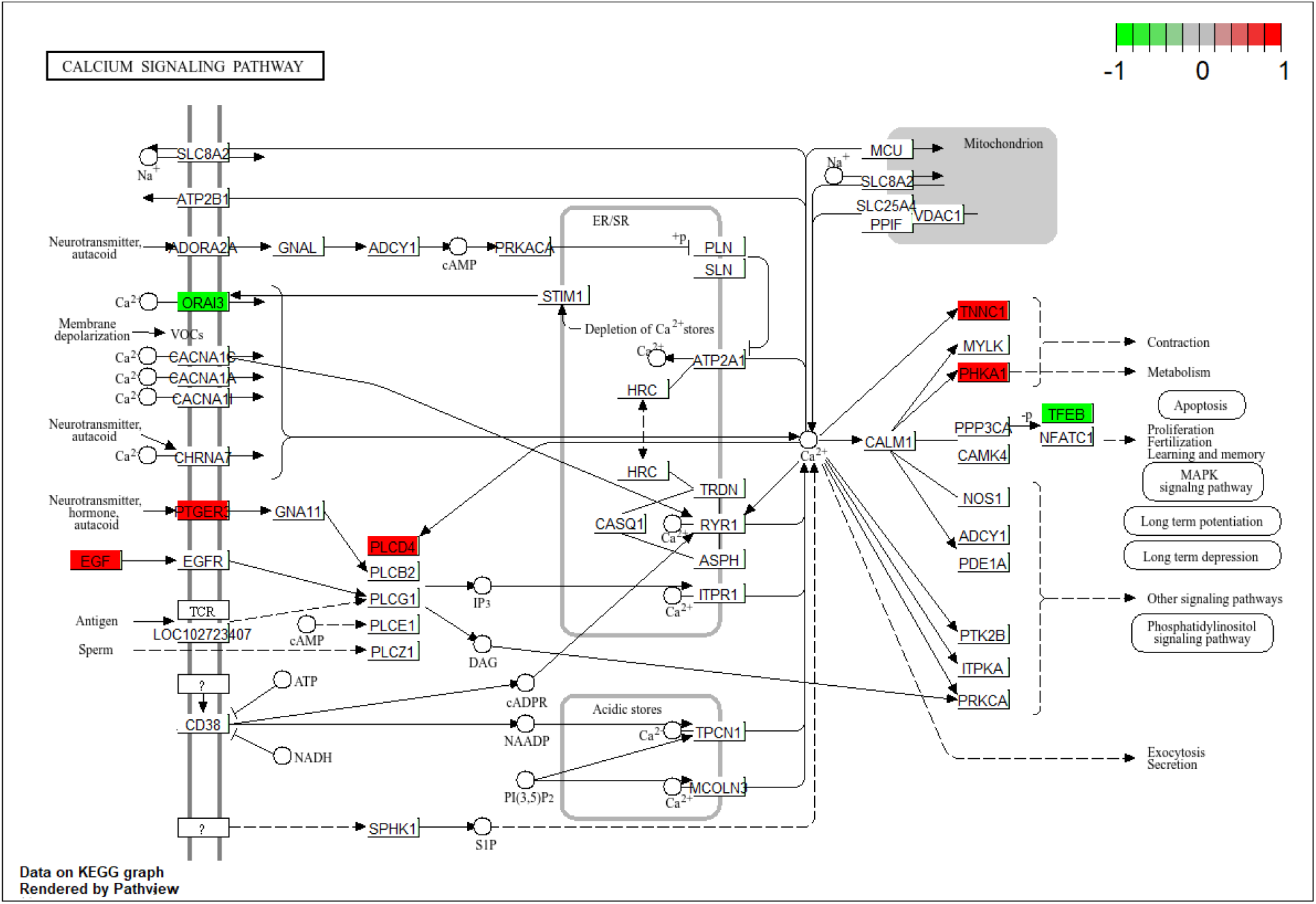
Crucial genes in the Calcium signaling pathways are up-regulated between HR and LR APOL1 variants in glomerular cells of patients with FSGS. The pathway chart depicts that EGF, PTGER3, PLCD4, TNNC1 and PHKA1 are up-regulated in the HR variants resulting in contraction – potentially of the glomerular tuft and vaso-constriction - whilst ORAI3 and TFEB are down-regulated.

Figure 4 shows a pathway chart of mTOR signaling highlighting genes up-regulated in APOL1 HR variants in red and genes down-regulated in green. The pathway chart depicts that the triggering WNT5A and TTI1 – one of the core genes in mTORC1 and mTORC2 – are up-regulated besides AMTOR3 and RRAGD, downstream of the Lysosome, and RPS6 involved in Ribosome biogenesis which Li et al. described to play a major role in FSGS via podocyte hypertrophy [48].

**Figure 4:**
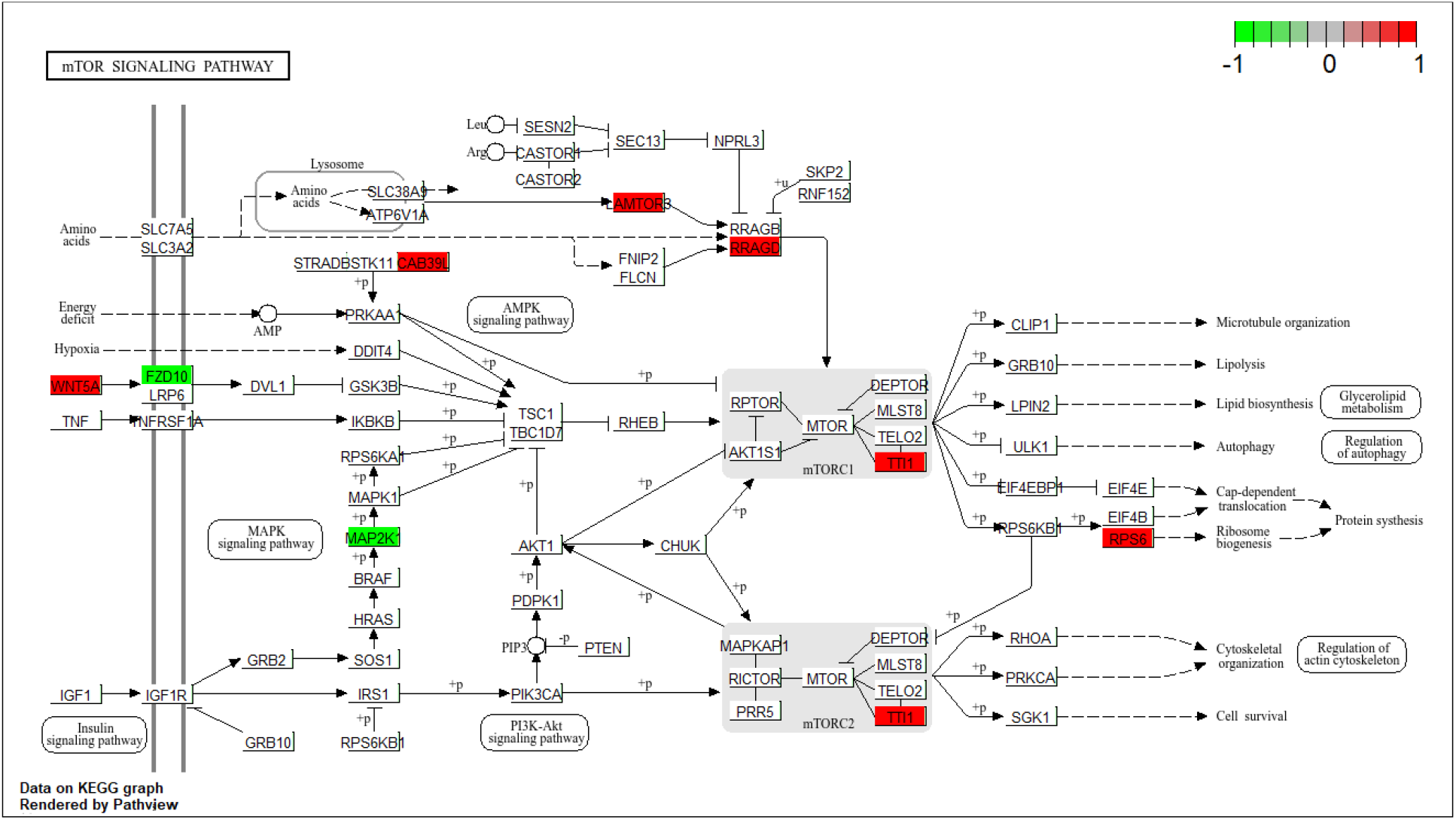
Crucial genes in the mTOR signaling pathway are up-regulated between HR and LR APOL1 variants. The pathway chart depicts that WNT5A and TTI1 - one of the core genes within the mTORC1 and mTORC2 complex– are up-regulated besides AMTOR3 and RRAGD, downstream of the Lysosome.Furthermore, we see upregulated expression of RPS6. Highlighting: red – up-regulated genes in APOL1 HR compared to LR variants, green – down-regulated genes.

### Gene-pathway networks operative in the glomeruli of FSGS HR patients

Figure 5 shows a gene-pathway network of genes up- and down-regulated in FSGS APOL1 HR versus LR variants in FSGS patient glomerular cells. Up-regulation denotes higher expression in APOL1 HR. The pathways in this network include our above-described findings of APOL1-variant-mediated differences in Calcium and mTOR signaling (Table 2, Figures 3 and 4). It has been proposed that inhibition of mTORc1 and APOL1 increase autophagy and reduce protein synthesis in the setting of chronic kidney disease [1], [49]. The inflammation-associated TNF signaling pathway appears to consist of predominantly down-regulated genes. Important hub genes are the down-regulated *MAP2K1* connecting mTOR signaling, TNF signaling and Long-term depression pathways. The up-regulated genes *GRIA3* and *TACR2* then connect the Long-term depression via Neuroactive ligand signaling with the Calcium signaling pathway. Neuroactive ligands signaling in podocytes again reflect the considerable overlap of pathways activated in brain and podocytes which we showed in an earlier publication [47]. RPS6 connects mTOR signaling to genes annotated under Coronavirus disease-COVID-19.

**Figure 5:**
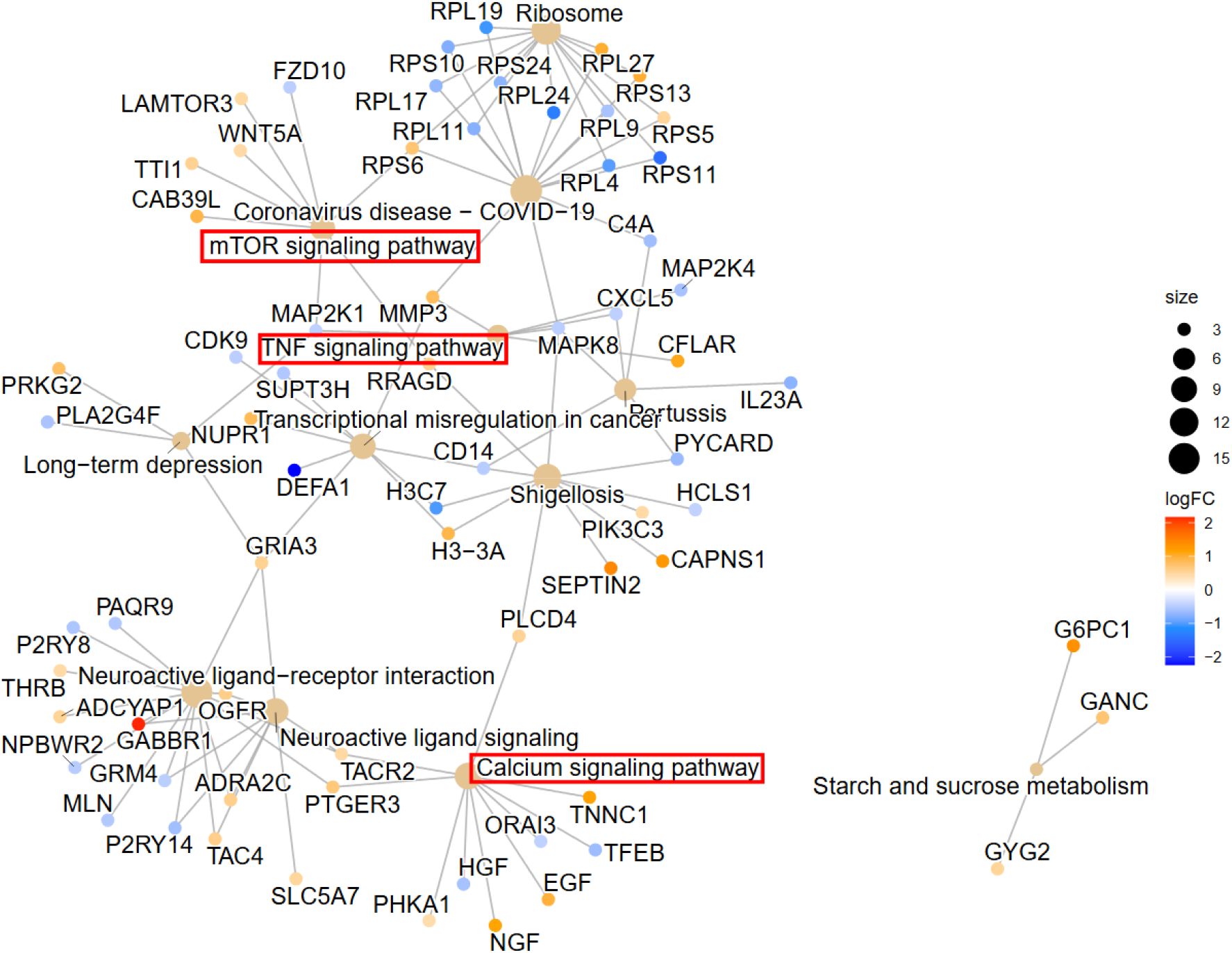
Gene-pathway network of genes significantly differentially expressed between APOL1 HR and LR variants in FSGS glomerular cells shows the involvement of Calcium and mTOR signaling pathways. Calcium and mTOR signaling pathways are affected by predominantly up-regulated genes in the HR variant while inflammation-related pathways such as TNF signaling are predominantly containing down-regulated genes.

### Confirmation of the FSGS glomerular biopsy-derived APOL1 HR gene signatures in in-vitro-models of APOL1 variant urine-derived podocytes

We set out to validate our gene signatures of up- and down-regulation in APOL1 HR from glomerular biospsies in FSGS patients from the NEPTUNE study (NCBI GEO accession GSE68125), [30]. The GEO dataset GSE194330 associated with the publication by Yoshida et al. [34] investigates gene expression of human urine-derived podocyte-like epithelial cells (HUPEC) from urine of APOL1 G0/G0 and G1/G2 genotypes. We compared significantly up- and down-regulated genes (p<0.05, fold change >1.5) in glomerular biopsies APOL1 HR vs. LR and HUPECs G1/G2 vs. G0/G0 and identified 25 genes down- and 31 genes up-regulated in common (Figure 6A, B). Over-representation analysis of Reactome pathways revealed Potassium channels in the 25 down-regulated genes and a plethora of pathways including starvation and stress response and mTOR signaling in the 31 up-regulated genes (Figure 6C, D, suppl. Table S4A, B). Moreover, we employed our gene signatures from the NEPTUNE study glomerular biopsies for heatmap and hierarchical cluster analysis in the APOL1 variant gene expression data in the HUPECs from Yoshida et al. [34]. Figure S2 depicts heatmaps and cluster analyses demonstrating that the gene signatures of up-regulated and down-regulated genes segregate the HUPEC APOL1 model in one G1/G2 and one G0/G0 cluster.

**Figure 6:**
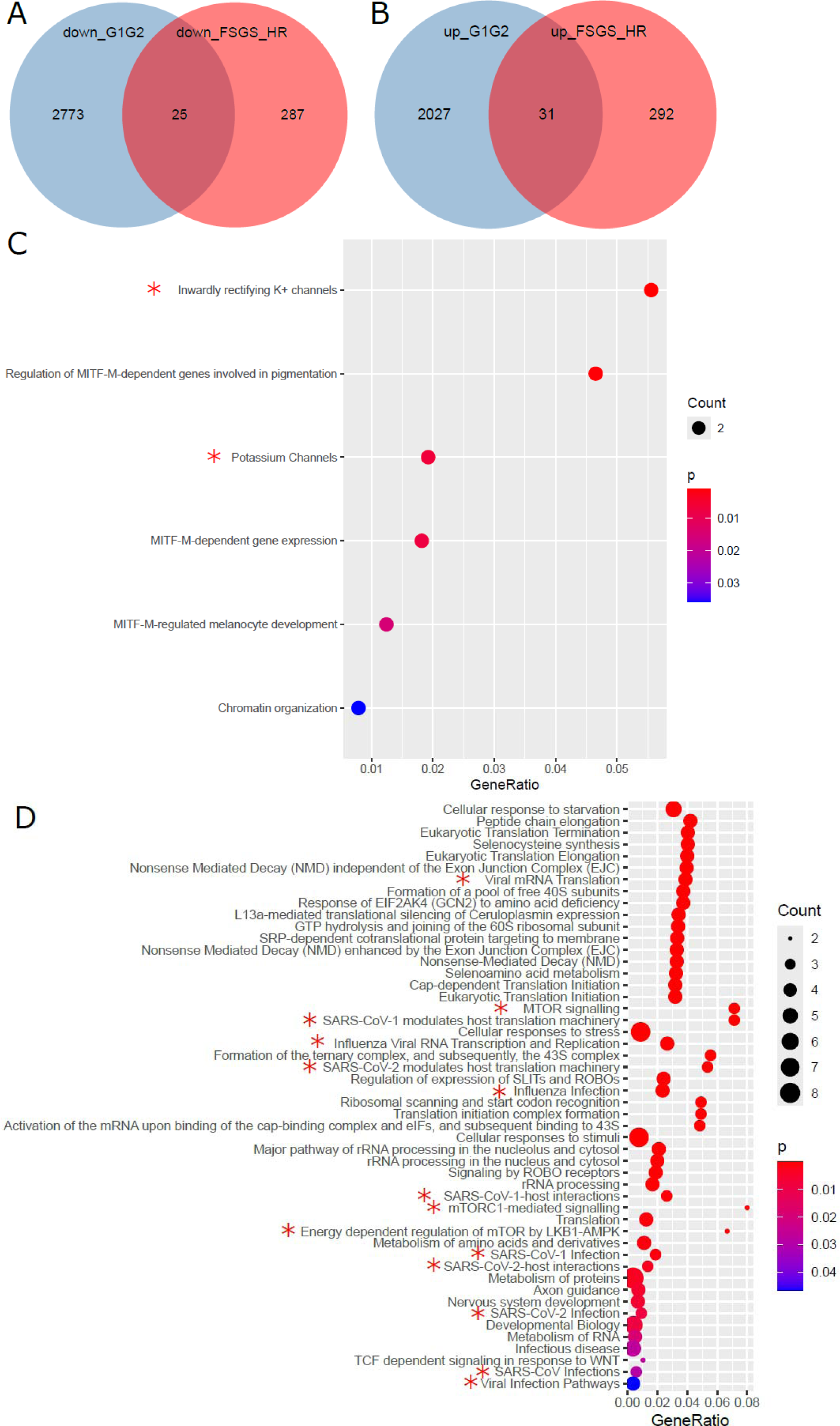
Overlap of up- and down-regulated genes in APOL1 HR variants between glomerular biopsies and HUPECs from the GEO dataset GSE194330. (A) Venn diagram analysis of genes down-regulated in APOL1 G1G2 variant HUPECs and APOL1 HR glomerular biopsies. (B) Venn diagram analysis of genes up-regulated in APOL1 G1G2 variant HUPECs and APOL1 HR glomerular biopsies. (C) Dot plot of significantly over-represented Reactome pathways in the 25 genes down-regulated in common in (A) including “Potassium channels”. (D) Dot plot of significantly over-represented Reactome pathways in the 31 genes up-regulated in common in (B) including “MTOR signaling” and several pathways related to viral infection.

The GEO dataset GSE85918 obtained from [9] is related with gene expression of human embryonic kidney cells (HEK293) with doxycycline-inducible (Tet-on) expression of APOL1 G0 and G1 genotypes.

We compared significantly up- and down-regulated genes (p<0.05, fold change >1.5) in glomerular biopsies APOL1 HR vs. LR and HEK293 Tet-on APOL1 G1 vs. G0 cells and identified 19 genes down- and 40 genes up-regulated in common (Figure 7A,B).

**Figure 7:**
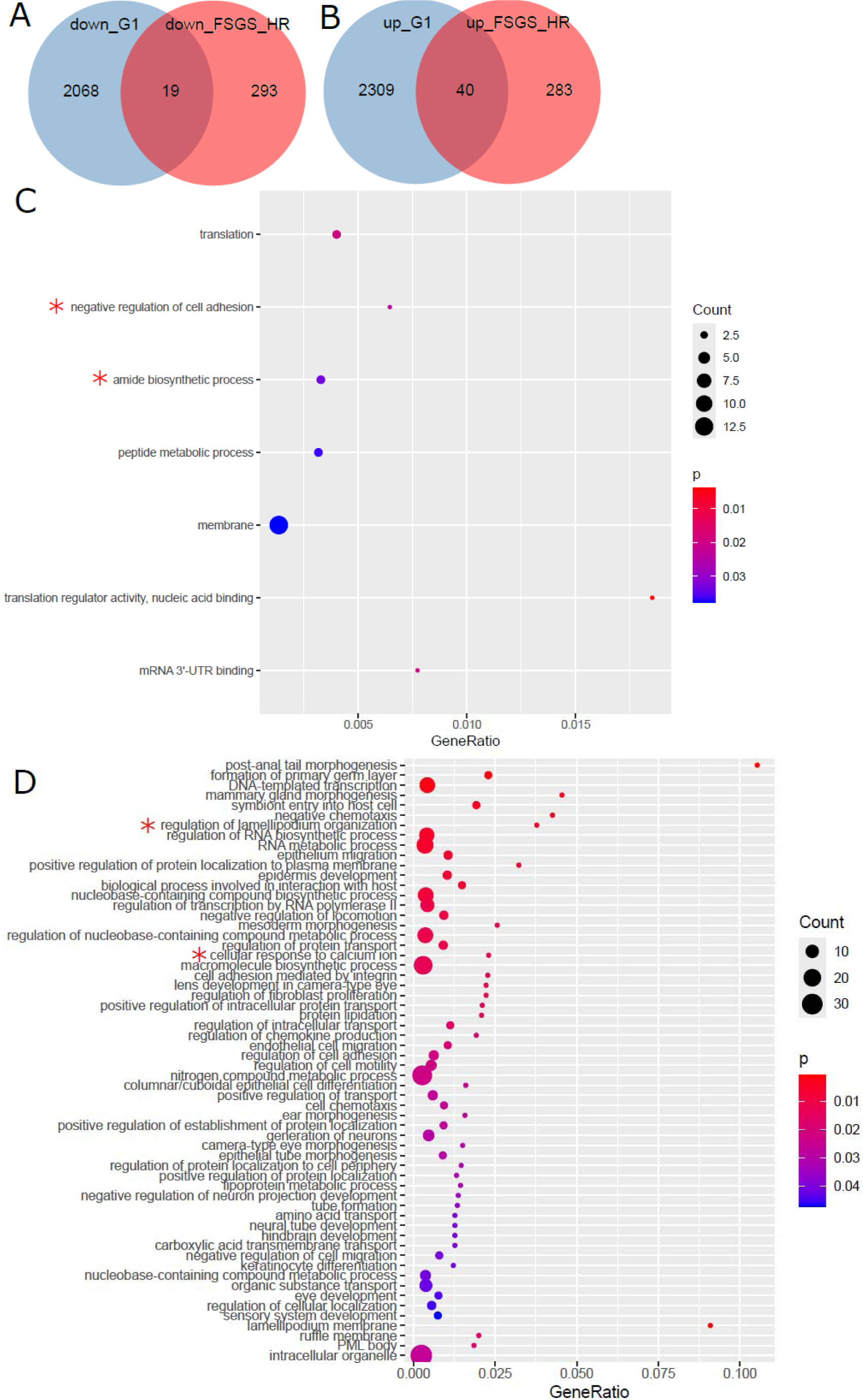
Overlap of up- and down-regulated genes in APOL1 HR variants between glomerular biopsies and HEK293 Tet-on APOL1 G0 and G1 cells from the GEO dataset GSE85918. (A) Venn diagram analysis of genes down-regulated in HEK293 Tet-on APOL1 G1 cells and APOL1 HR glomerular biopsies. (B) Venn diagram analysis of genes up-regulated in HEK293 Tet-on APOL1 G1 cells and APOL1 HR glomerular biopsies. (C) Dot plot of significantly over-represented Gene ontologies (GOs) in the 19 genes down-regulated in common in (A) including “negative regulation of cell adhesion” and “amide biosynthetic process”. (D) Dot plot of significantly over-represented GOs in the 40 genes up-regulated in common in (B) including “regulation of lamellipodium organization” and “cellular response to calcium ion”.

The dot plot of significantly over-represented Gene ontologies (GOs) in the 19 genes down-regulated in common (Figure 7A) in Figure 7C shows terms including “negative regulation of cell adhesion” and “amide biosynthetic process”. The dot plot (Figure 7D) of significantly over-represented GOs in the 40 genes up-regulated in common in (B) includes “regulation of lamellipodium organization” and “cellular response to calcium ion”.

Furthermore, we employed our gene signatures from the NEPTUNE study glomerular biopsies for heatmap and hierarchical cluster analysis in the APOL1 variant gene expression data in the HEK293 cells [9]. Figure S3 depicts heatmaps and cluster analyses demonstrating that the gene signatures of up-regulated and down-regulated genes also segregates the HEK293 cells into a G1 and G0 cluster respectively

The GEO dataset GSE135663 from [33] investigates single-cell RNAseq of kidney organoids with genome-edited G1 genotypes stimulated with IFN-γ. We employed our gene signatures from the NEPTUNE study glomerular biopsies for heatmap and hierarchical cluster analysis in the APOL1 variant gene expression data in the kidney organoids from Liu et al. [33]. Figure S3 depicts heatmaps and cluster analyses distinct gene clusters up-regulated and down-regulated in the G1 genotype and IFN-γ stimulation. The heatmap in Figure S3A uses the signature of genes up-regulated between APOL1 HR and LR in the glomerular biopsies to show the mean gene expression over all single cells for the APOL1 G1 and G0 kidney organoids stimulated with IFN-γ. As APOL1 is activated by IFN-γ gene clusters up-regulated between G1-IFN (red color bar) and G0-IFN (blue color bar) are regulated in the same direction as in the biopsy data. Figure S3B shows the analogous heatmap for genes down-regulated between APOL1 HR and APOL1 LR in the glomerular biopsies.

### Identification of small molecules reverting transcriptome chances induced by APOL1 HR variants

We set out to investigate via the webtool SigCom LINCS if there are small molecules (SMs) which are capable of reverting the transcriptomic changes induced by the APOL1 HR variants. Figure 8 shows that the mTOR inhibitor AZD-2014 *in silico* reverses the signature of genes up- and down-regulated by IFN-γ -stimulation in APOL1 G1-variant kidney. Figure 8A depicts a SigCom LINCS heatmap of small molecules reverting (orange) and mimicking (violet) scRNAseq gene expression of G1-variant kidney after IFN-γ stimulation. The gene set is successively refined to the top regulated genes in Figure 8B and to cluster1 including the most consistently down-regulated ranks (Figure 8C). The genes from the down-regulated cluster and their gene expression ranks in the SM signatures are provided as Supplementary Table S6. Several small molecules reverse the IFN-γ stimulated gene expression to the non-stimulated gene expression. These include in the human kidney cell line HA1E, BRD-K44573794 (PubChem CID: 54652754; IUPAC Name: (1R,2aR,8bR)-2-(1,3-benzodioxol-5-ylmethyl)-1-(hydroxymethyl)-N-propyl-1,2a,3,8b-tetrahydroazeto[2,3-c]quinoline-4-carboxamide), BRD-K26169592 (PubChem CID: 44488019; IUPAC Name: N-[(2S,3S)-5-[(2S)-1-hydroxypropan-2-yl]-3-methyl-2-[[methyl(naphthalen-1-ylcarbamoyl)amino]methyl]-6-oxo-3,4-dihydro-2H-1,5-benzoxazocin-10-yl]pyridine-4-carboxamide), BRD-K67548518 (PubChem CID: 73819568; IUPAC Name: N-[(2S,4aS,12aR)-2-[2-(2,3-dihydro-1H-inden-2-ylamino)-2-oxoethyl]-5-methyl-6-oxo-2,3,4,4a,12,12a-hexahydropyrano[2,3-c][1,5]benzoxazocin-8-yl]-4-(trifluoromethyl)benzamide) and the mTOR inhibitor-AZD-2014 (highlighted in green). EnrichR enrichment analysis of the genes from cluster1 yielded results shown in Figures 8D-F including *IgA Nephropathy* from EnrichR dataset GWAS_Catalog_2025 (Figure 8D), *Interferon signaling* (Figure 8E) from EnrichR dataset Reactome pathways and *Lumenal side of Endoplasmic reticulum membrane* and *Phagocytic vesicle* (Figure 8F) from EnrichR dataset Gene Ontology – Cellular component.

**Figure 8:**
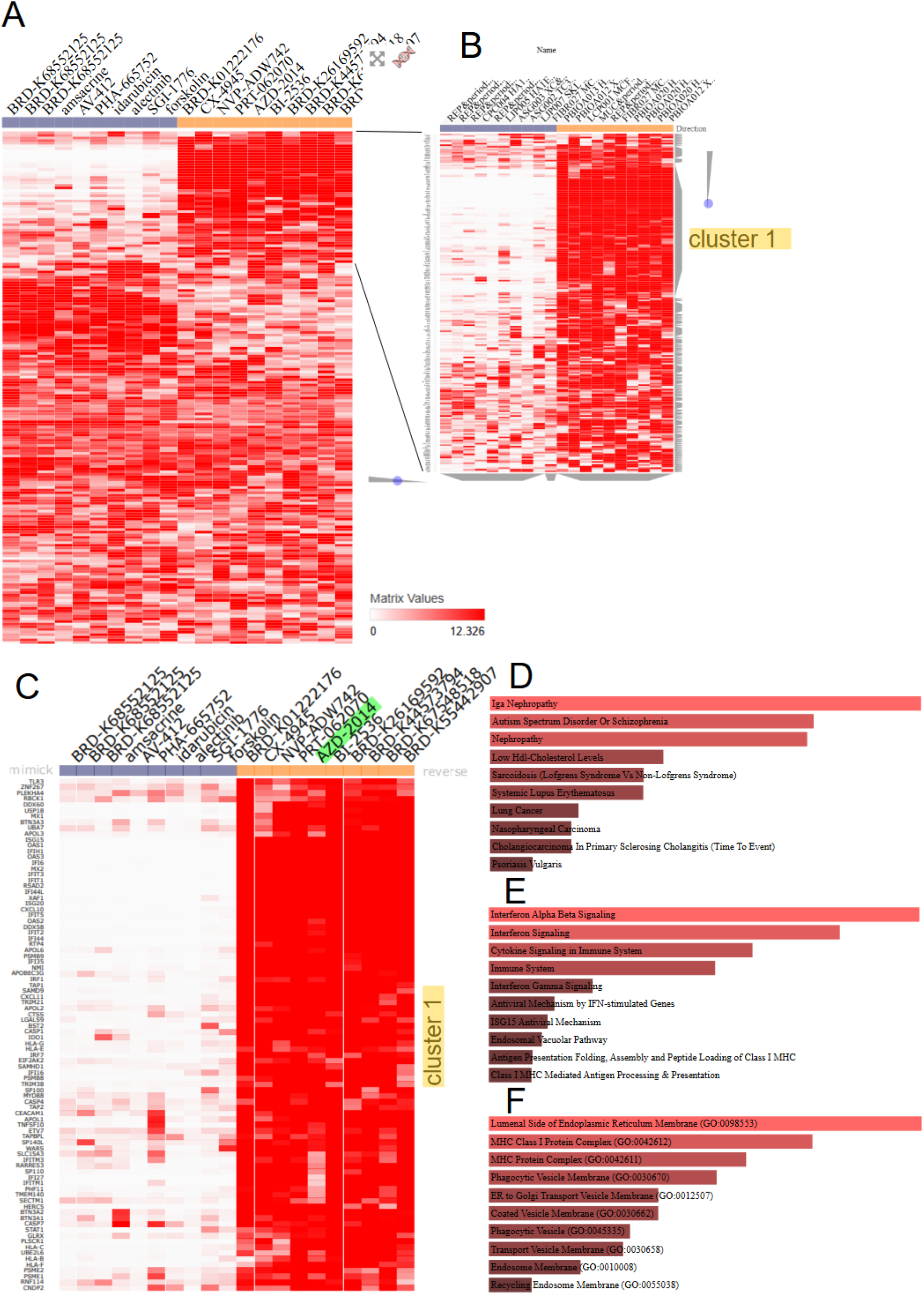
The mTOR inhibitor AZD-2014 *in silico* reverses the signature of genes up- and down-regulated by IFN-γ -stimulation in APOL1 G1-variant iPSC-derived kidney organoids. (A) SigCom LINCS Heatmap of small molecules reverting (orange) and mimicking (violet) scRNAseq gene expression of G1-variant kidney after IFN-γ stimulation. The gene set is step-wise refined to the top regulated genes in (B) and to cluster1 including the most consistently down-regulated ranks (C). Several small molecules reverse the regulation after IFN-γ stimulation, these include in the kidney cell line HA1E - BRD-K44573794, BRD-K26169592 and BRD-K67548518 and the mTOR inhibitor AZD-2014 (highlighted in green). The genes from clsuter1 were subjected to EnrichR enrichment analysis. (D) The bar chart shows the most significantly enriched results from EnrichR dataset GWAS_Catalog_2025 with *IgA Nephropathy* on top. (E) The bar chart shows the most significantly enriched results from EnrichR dataset Reactome pathways which includes *Interferon signaling*. (F) The bar chart shows the most significantly enriched results from EnrichR dataset Gene Ontology – Cellular component including *Lumenal side of Endoplasmic reticulum membrane* and *Phagocytic vesicle*.

## Discussion

In this meta-analysis, we compared gene expression in APOL1 HR and LR variants after IFN-γ stimulation in kidney biopsies, organoids and cell lines. In the kidney biopsies obtained from the NEPTUNE study [30] we focused on glomerular cells of patients with FSGS and identified signatures of genes up- and down-regulated between APOL1 HR and LR variants. The gene signature derived from the biopsy data enabled segregation of APOL1 HR and LR variant associated transcriptomes in the cellular models.

Furthermore, venn-diagram based analyses revealed overlapping intersection sets of 25 down-and 31 up-regulated genes between the HUPEC model and biopsy data, and 19 down-regulated and 40 up-regulated genes between the HEK293 model and biopsy data. Considering over-represented gene ontologies, the HUPEC model revealed potassium channels and mTOR signaling whilst the HEK293 model identified more general terms such as translation in the down-regulated but at least some more specific terms such as “Cellular response to calcium ion” in the up-regulated genes.

In the up-regulated genes we identified the Calcium and mechanistic target of rapamycin (mTOR) signaling pathways over-represented whilst in the down-regulated genes the results include inflammation-associated pathways such as Toll-like receptor and TNF signaling. Two sources of cellular Calcium cations (Ca^2+^) exist: (i) extracellular Ca^2+^ can flow into the cell via cation pores and (ii) Ca^2+^ from the endoplasmic reticulum (ER) can flow into the cytosol via channels such as the inositol trisphosphate receptors (IP3Rs) and ryanodine receptors (RYRs). While Datta et al. propose Ca^2+^ release from the ER via RYR and IP3R [1], Adebayo et al. report increased Ca^2+^ influx through plasma membrane pores in case of APOL1 HR variants and show that this is the main source of Ca^2+^ leading to cytotoxicity [50]. Furthermore, the protective APOL1 variant N264K (M1) [50] [51] and the APOL1 channel blocker VX147 can inhibit Ca^2+^ influx [50]. mTOR signaling regulates fundamental cellular processes such as cell growth, protein synthesis, metabolism, proliferation and autophagy and its de-regulation is involved in aging, diabetes and cancer [52]. mTOR signaling maintains renal cell homeostasis and in acute kidney injury (AKI) can foster kidney regeneration while constant mTOR activation can result in glomerular hypertrophy, interstitial fibrosis, polycystic kidney disease, and renal cell carcinoma [53]. Efficiency of mTOR inhibitors in particular rapamycin has been demonstrated in several studies for many renal diseases [53], [54], also including kidney fibrosis [55]. mTOR inhibition increases autophagy and reduces protein synthesis. Surprisingly, Datta et al. showed that inhibition of mTOR leads to a downstream cascade of events following APOL1-G1-mediated cation pore formation in the plasma membrane [1]. The authors report G1-mediated Na+ import/K+ efflux triggered activation of GPCR/IP3–mediated calcium release from the ER, impaired mitochondrial ATP production, phosphorylation (activation) of AMPK by CaMKKβ promoting autophagy and inhibiting protein synthesis, via inhibition of mTORC1 and eIF2α, which were all reversed by the APOL1 channel blocker VX-147 [1]. The inhibition of mTOR in this setting is to some extent surprising as others have reported activated mTOR in FSGS [48], [56] and CKD [53] - in line with our in silico modeling results of up-regulated mTOR signaling in HR vs. LR APOL1-mediated FSGS and efficiency of the mTOR inhibitor AZ-124 in reversing the APOL1 G1 signature after IFN-γ -stimulation. The observation of inhibited mTOR may be a short-term response measured after 8h of Tet-induced APOL1-G1 variant in the transformed T-REx-293 G1 cell line *in vitro* and the up-regulated mTOR in kidney biopsy-derived data may reflect differences between *in vivo* long-term effects and *in vitro* glomerular biopsy-derived cells. However, inhibition and persistent up-regulation of mTOR, can both have detrimental effects on podocytes and rapamycin treatment can lead to proteinuria, FSGS and impairment of renal function in the context of existing glomerular disease [56], [57], [58].

We conclude that our study has revealed a signature of genes up- and down-regulated in glomerular cells of NEPTUNE-FSGS patients expressing the APOL1 HR variant. Genes up-regulated in APOL1-HR were associated with Calcium and mTOR signaling. The gene signature was able to segregate HR from LR samples in cellular AMKD models and to some extent up- and down-regulated genes overlapped. When associating the overlapping genes with biological terms, the patient-specific HUPEC (Human urine-derived podocyte epithelial cells) model was much closer to the NEPTUNE-FSGS biopsy data than the HEK293 model with regards to mTOR signaling. Analysis of small molecules reverting the IFN-γ stimulated single-cell gene expression to the non-stimulated gene expression in genome-edited APOL-G1 kidney organoids revealed several putative drug candidates, amongst them the mTOR inhibitor-AZD-2014.

## Declarations

### Ethics approval and consent to participate

Not applicable.

### Consent for publication

Not applicable

### Availability of data and materials

The datasets analyzed during the current study are available in the National Center for Biotechnology Information (NCBI) Gene expression Omnibus (GEO) repository, accessions GSE279611 (https://www.ncbi.nlm.nih.gov/gds/?term=GSE279611), GSE171240 (https://www.ncbi.nlm.nih.gov/gds/?term=GSE171240) and GSE186823 (https://www.ncbi.nlm.nih.gov/gds/?term=GSE186823.

An overview of the datasets used is provided in Table 1.

### Contributions

JA initiated the concept for this study. WW and CT analysed the data and wrote the manuscript. JA supervised the work, co-wrote the manuscript and gave the final approval. TA, MDSS and SP provided technical support.

## Acknowledgments

James Adjaye acknowledges financial support from the Medical faculty of the Heinrich-Heine University, Duesseldorf.

## Supplementary Material

**Supplementary Information (Supplementary_information_wruck_etal.pdf)**

**Supplementary Table S1. (tableS1.xlsx): In vitro cellular models for APOL1 mediated kidney disease.**

**Supplementary Table S2. (tableS2.xlsx): Characteristics of glomerular cell samples of FSGS patients from the NEPTUNE project.**

**Supplementary Table S3. (tableS3.xlsx): significantly up- and down-regulated genes between APOL1-HR vs -LR variants in the glomerular cells from FSGS patients of the NEPTUNE project.**

**Supplementary Table S4. (tableS4.xlsx): significantly over-represented Reactome pathways.**

**Supplementary Table S5. (tableS5.xlsx): Kidney cell line HA1E statistics of small molecules reverting IFN-γ -induced APOL1-HR-variant-mediated gene expression.**

**Supplementary Table S6. (tableS6.xlsx): up- and down-regulated genes in IFN-γ -induced APOL1-HR-variant-mediated gene expression, their ranks in SigCom LINCS SM analysis and enrichment analysis results.**

## Supplementary Figures

**Figure S1:**
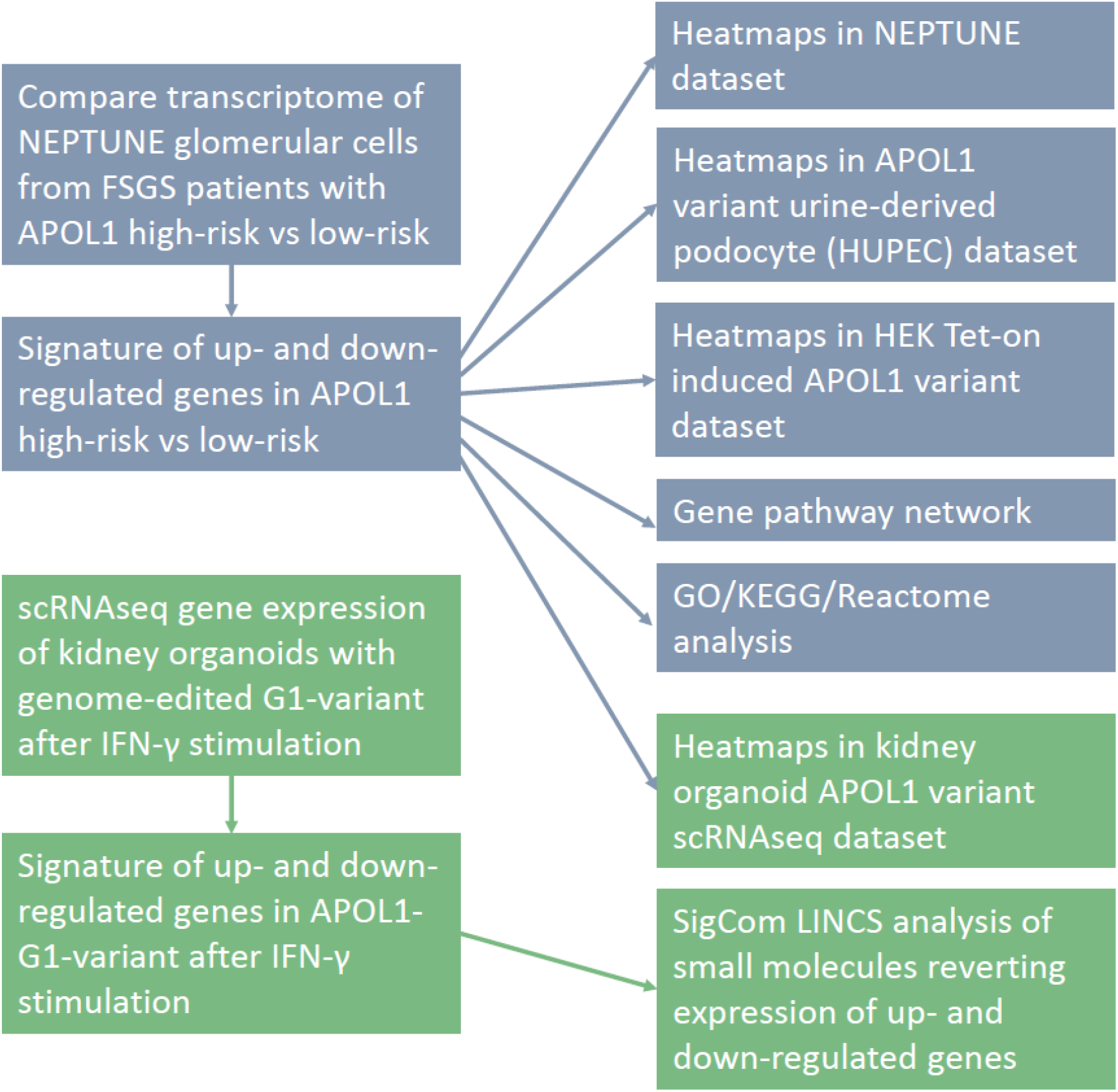
Pipeline of analyses performed in the NEPTUN biopsy data, the urine-derived podocytes, the HEK model and the single-cell-RNAseq of kidney organoids.

**Figure S2:**
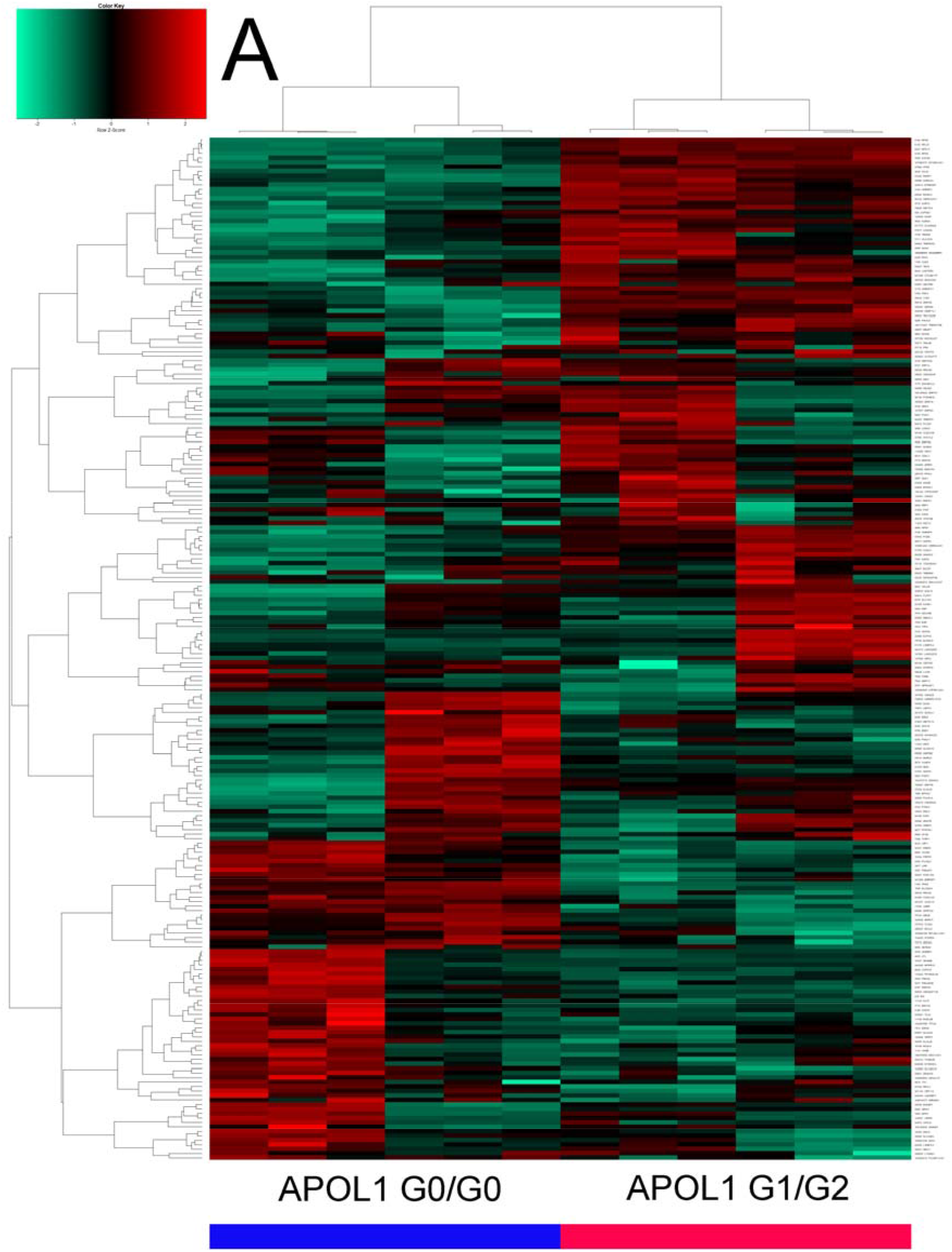
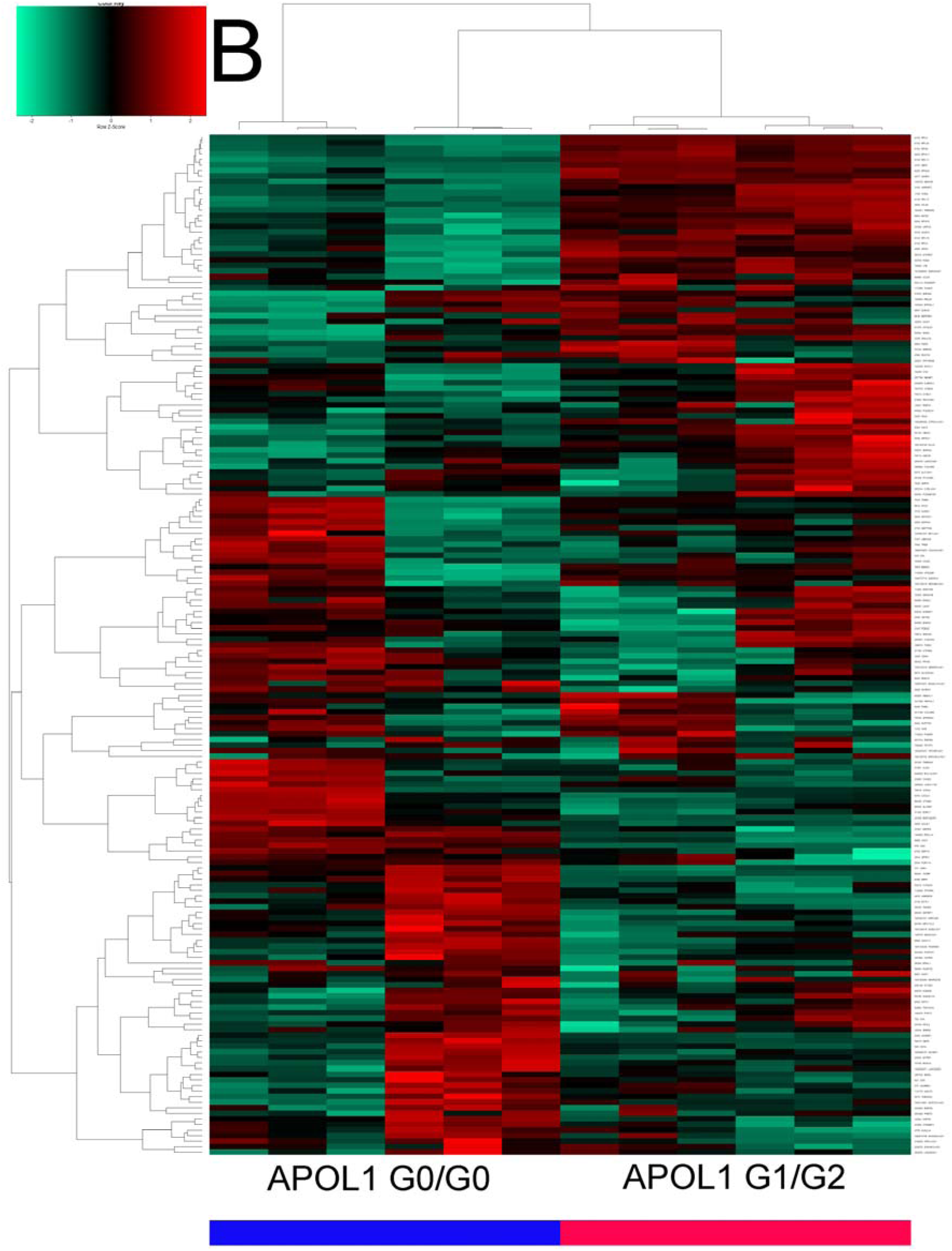
Gene signatures from the APOL1 HR glomerular biopsies can distinguish between APOL1 G1/G2 and G0/G0 individuals in HUPECs from the GEO dataset GSE194330. (A) Genes up-regulated between APOL1 HR and LR in the glomerular biopsies segregates HUPEC samples into a G1/G2 and G0/G0 cluster respectively. (B) Genes down-regulated between APOL1 HR and APOL1 LR in the glomerular biopsies segregates HUPEC samples into a G1/G2 and G0/G0 cluster. (color bar: blue – APOL1 G0/G0 samples, red – APOL1 G1/G2 samples).

**Figure S3:**
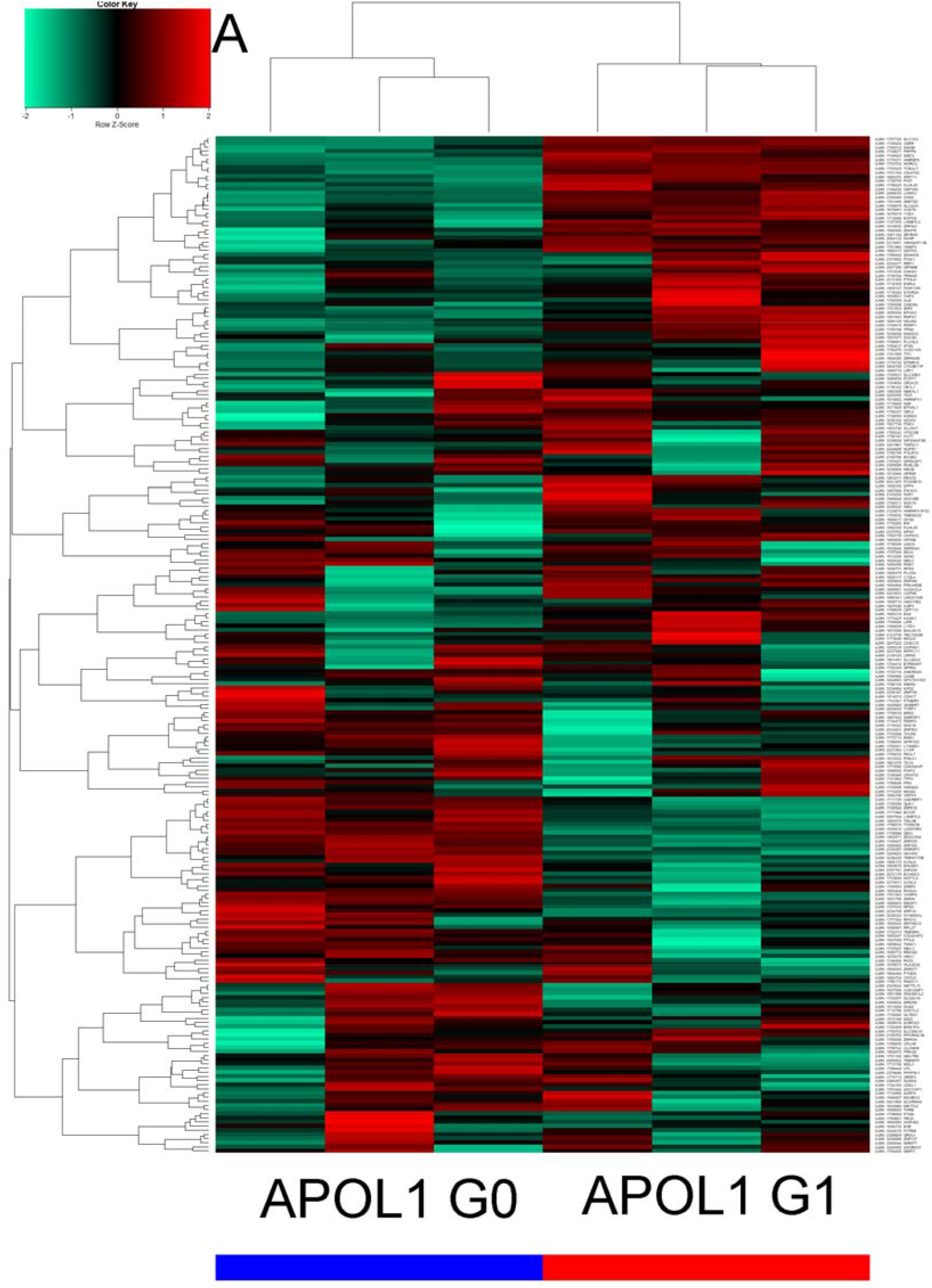
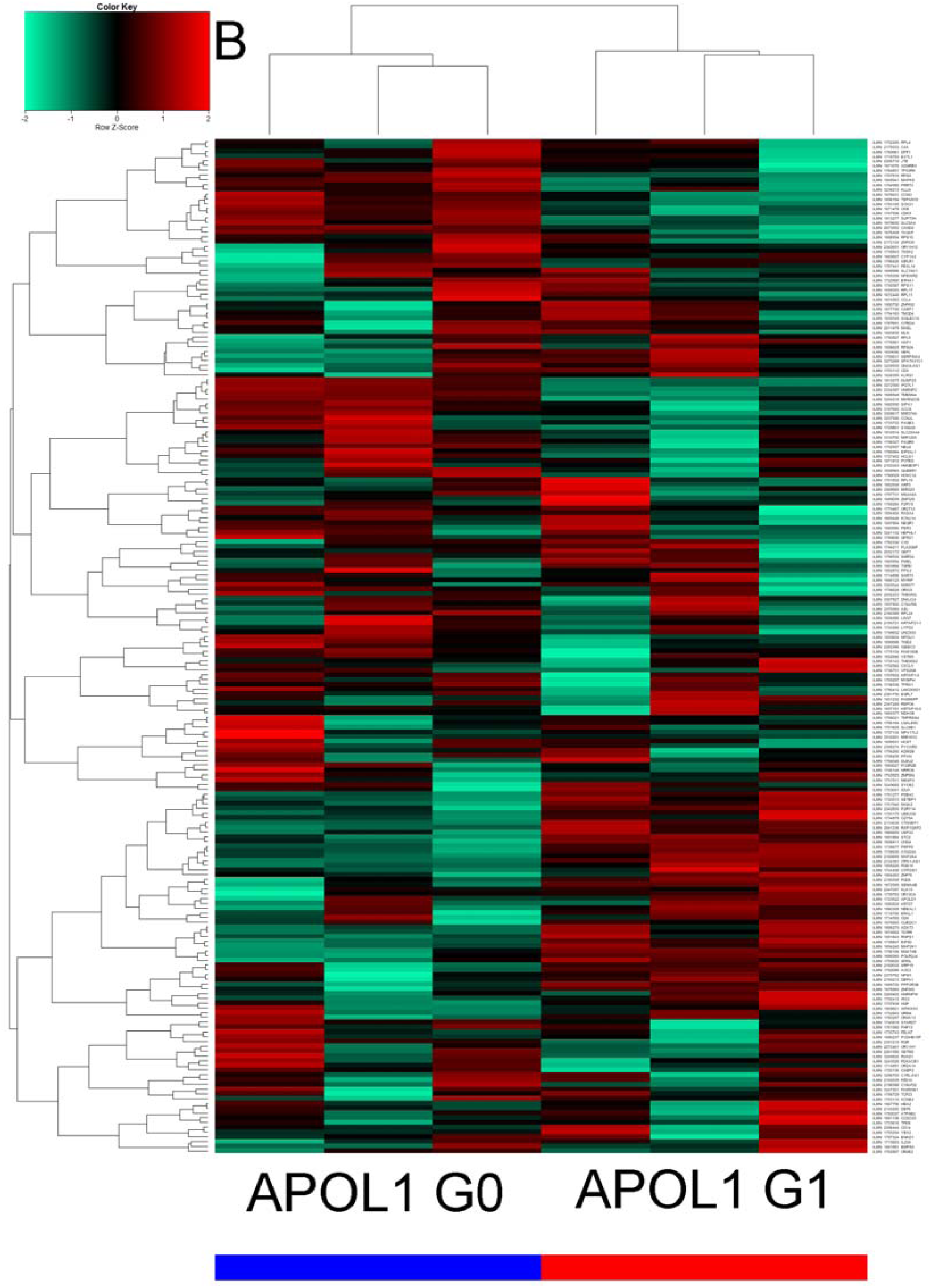
Gene signatures from the APOL1 HR glomerular biopsies can distinguish between APOL1 G1 and G0 individuals in HEK293 Tet-on APOL1 G0 and G1 cells from the GEO dataset GSE85918. (A) Genes up-regulated between APOL1 HR and LR in the glomerular biopsies separate HEK293 Tet-on APOL1 G0 and G1 samples into one G1 and one G0 cluster. (B) Additionally, genes down-regulated between APOL1 HR and APOL1 LR segregate HEK293 Tet-on APOL1 G0 and G1 samples into one G1 and one G0 cluster. (color bar: red – APOL1 G1 samples, blue – APOL1 G0 samples).

**Figure S4.**
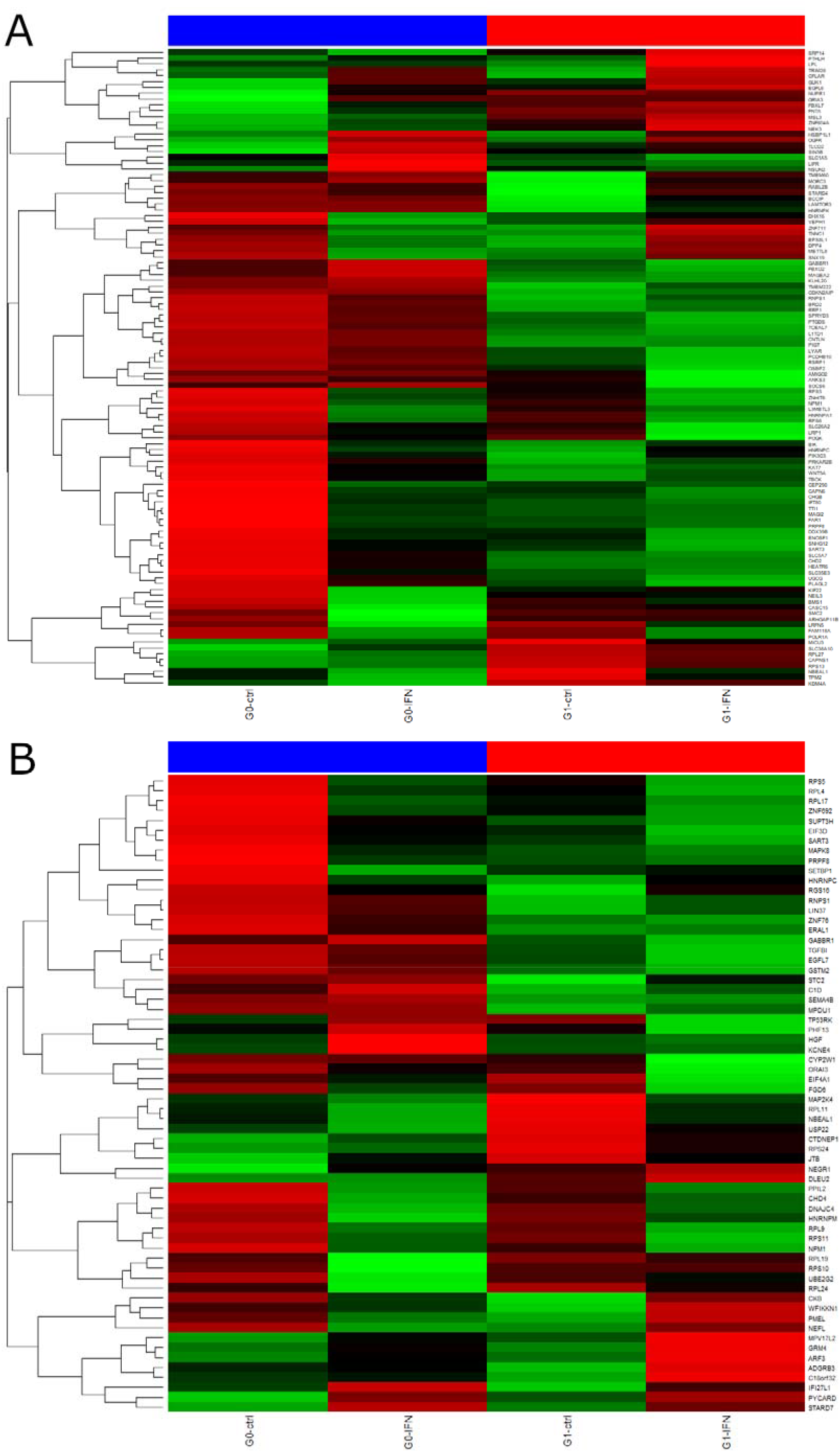
(figureS4.png): Gene signatures from the APOL1 HR glomerular biopsies were employed to investigate gene expression in single-cell RNA-seq experiments between APOL1 G1 and G0 iPSC-derived kidney organoids stimulated with IFN-γ from the GEO dataset GSE135663. (A) Genes up-regulated between APOL1 HR and LR in the glomerular biopsies identify gene clusters up- and down-regulated between APOL1 G1 and G0 kidney organoids stimulated with IFN-γ. (B) Additionally, genes down-regulated between APOL1 HR and APOL1 LR in the glomerular biopsies identify gene clusters up- and down-regulated between APOL1 G1 and G0 iPSC-derived kidney organoids stimulated with IFN-γ. (color bar: red – APOL1 G1 samples, blue – APOL1 G0 samples).

