## Supplementary Information for "Meta-analyses of the NEPTUNE dataset related to high and low risk FSGS and cellular APOL1 models identifies Calcium signalling, mTOR signaling and inflammation-associated pathways as operative in APOL1-mediated kidney disease"

### Supplementary information for "A Meta-Analysis of APOL1 risk variants in single-cell and bulk transcriptome data from kidney organoids and biopsies." by Wruck et al.

##### Overview of findings from in vitro cellular models for APOL1 mediated kidney disease

Table S1 provides an overview of in vitro cellular models for APOL1 mediated kidney disease. Khatus et al. found with their HEK-based model that Exon 4 determines APOL1 cytotoxicity (Khatua et al., 2015). Zimmermann et al. tested Inaxiplin as an potenitial APOl1 channel inhibitor in vitro and in vivo (Zimmerman et al., 2025). Chun et al. reported that (i) APOL1 in G0, is mostly localised in lipid droplets (LDs), whereas the risk variants G1 and G2 reside mainly in the endoplasmic reticulum (ER). This influences cellular toxicity, with ER retention of the risk variants contributing to cell damage, and that (ii) LD recruitment of the risk variants can allow for protection against cell toxicity and autophagic flux impairment (Chun et al., 2019) . Xiao et al. via RNA-seq data revealed (i) that HIF2α can regulate APOL1 and lncRNA LINC02609 expression and found that HIF-2α can bind to the promoter of APOL1 and lncRNA LINC02609 and transcriptionally regulate their expression directly, and (ii) showed that high APOL1 expression correlated with worse clinical outcomes, and knockdown of APOL1 inhibited tumor cell lipid droplet formation, proliferation, metastasis and xenograft tumor formation abilities (Xiao et al., 2024). Untreated HUVECs were compared to IFNɣ-exposed; and APOL1 expression, mitochondrial function, lysosome integrity, and autophagic flux were measured (Blazer et al., 2022). IFNɣ increased median APOL1 expression across all genotypes 22.1 (8.3 to 29.8) fold ($p=0.02$). Compared to zero risk variant-carrying HUVECs (0RV), HUVECs carrying 2 risk variant copies (2RV) showed both depressed baseline and maximum mitochondrial oxygen consumption ($p<0.01$), and impaired mitochondrial networking on MitoTracker assays. These cells also demonstrated a contracted lysosomal compartment, and an accumulation of autophagosomes suggesting a defect in autophagic flux. UM30-OSN cells represent a valuable additional model for investigating kidney-associated diseases such the contribution of APOL1 high-risk variants to kidney injury and fibrosis (Thimm et al., 2025). Wakashin et al. (i) confirmed mRNA splice variants in the kidney and those who contribute to kidney disease via TA-cloning. Six mRNA variants found: V1 (A), V3 (A), V2–1 (B1) and V4 (C);APOL1 isoform APOL1-B3 regulates inflammatory signaling and reported (ii) that APOL1-B3, both wild- type (G0) and renal risk variant (G2), enhanced the early events of IL-1 receptor signaling, the phosphorylation of MAPK and ERK in APOL1-B3-HeLa cells (Wakashin et al., 2020). Ekulu et al. induced APOL1 through polyinosinic-polycytidylic acid (poly(I:C)) what caused podocyte detachment, decline in cell viabiltiy and increased apoptosis rate in a genotype independent manner upregulation of CD2AP, alteration of cytoskeleton, reduction of autophagic flux and increased permeability in an in vitro model under continuous perfusion (Ekulu et al., 2021).G2/G2 podocyte showed altered features which include up. Ma et al. reported (i) that in human kidney, APOL1 is derived from both endogenous synthesis and extracellular sources, which also include protein uptake, (ii) that podocytes show higher levels of APOL1 protein in kidney tissues due to both local synthesis and uptake from circulation or glomerular filtrate, (iii) that in vitro, exogenous APOl1 can actively be taken up by podocytes, which suggest that circulating APOl1 may contribute to its renal accumulation and (iv) that APOL1 mRNA is present in many different kidney cell types (Ma et al., 2015). Carracedo et al. show that (i) APOL1 high-risk variants (G1, G2) induce endothelial activation, marked by increased ICAM1 and reduced PECAM1 expression in iPSC-derived and primary glomerular endothelial cells, that (ii) Endothelial activation is cell-autonomous, leading to enhanced monocyte adhesion independent of external inflammatory stimuli and that (iii) APOL1 expression drives endothelial dysfunction beyond the glomerulus, supporting a role for APOL1 risk variants in inflammatory renal angiopathy (Carracedo et al., 2023). Via single-cell transcriptomic analysis and immunofluorescence images Song et al. showed (i) upregulation of APOL1 in podocytes after interferon-gamma treatment, (ii) significant reduction was seen in oxidative phosphorylation and TCA (tricarboxylic acid) cycle activity; glycolysis and hypoxia signaling in risk variants podocytes were upregulated, (iii) the isolated risk variant glomeruli showed no increase in respiration rate after IFN-gamma treatment, while the iPSC-derived RV podocytes showed reduced mitochondrial branches and shorter branch length (Song et al., 2025).

Juliar et al. report that (i) exposure to IFN-gamma degrades endothelial networks in kidney organoids, that (ii) APOL1 and pyroptosis -associated genes were upregulated but not apoptosis related genes, that (iii) endothelial networks were rescued when JAK/STAT signaling or pan-caspase were inhibited and that (iv) patients with accelerated renal failure had upregulated IFN and pyroptosis genes (Juliar et al., 2024). Song et al. also (i) established an APOL1-associated kidney model from G1 and G2 patients using iPSC-organoids, (ii) found that APOL genetic risk variants affect population of podocyte and (iii) showed in their analysis metabolic reprogramming in risk variant podocytes as well as decline in functional and structural mitochondrial function (Song et al., 2025).

#### Supplementary tables

**Table S1 (tableS1.xlsx): In vitro cellular models for APOL1 mediated kidney disease.**

**Supplementary Table S2 (tableS2.xlsx): Characteristics of glomerular cell samples of FSGS patients from the NEPTUNE project.**

**Supplementary Table S3 (tableS3.xlsx): significantly up- and down-regulated genes between APOL1-HR vs -LR variants in the glomerular cells from FSGS patients of the NEPTUNE project.**

**Supplementary Table S4 (tableS4.xlsx): significantly over-represented Reactome pathways.**

**Supplementary Table S5 (tableS5.xlsx): Kidney cell line HA1E statistics of small molecules reverting IFN-γ -induced APOL1-HR-variant-mediated gene expression.**

**Supplementary Table S6 (tableS6.xlsx): up- and down-regulated genes in IFN-γ -induced APOL1-HR-variant-mediated gene expression, their ranks in SigCom LINCS SM analysis and enrichment analysis results.**
